# Sample preparation methods for volume electron microscopy in mollusc *Berghia stephanieae*

**DOI:** 10.1101/2024.02.25.581936

**Authors:** Brandon Drescher, Harshada H. Sant, Richard L. Schalek, Jeff W. Lichtman, Paul S. Katz

## Abstract

Creating a high-resolution brain atlas in diverse species offers crucial insights into general principles underlying brain function and development. A volume electron microscopy approach to generate such neural maps has been gaining importance due to advances in imaging, data storage capabilities, and data analysis protocols. Sample preparation remains challenging and is a crucial step to accelerate the imaging and data processing pipeline. Here, we introduce several replicable methods for processing the brains of the gastropod mollusc, *Berghia stephanieae* for volume electron microscopy. Although high-pressure freezing is the most reliable method, the depth of cryopreservation is a severe limitation for large tissue samples. We introduce a BROPA-based method using pyrogallol and methods to rapidly process samples that can save hours at the bench. This is the first report on sample preparation and imaging pipeline for volume electron microscopy in a gastropod mollusc, opening up the potential for connectomic analysis and comparisons with other phyla.

## Introduction

Growing interest in volume electron microscopy requires technological development to accommodate serial sectioning of large sample volumes (Schalek et al., 2012), imaging thousands of acquired sections (Denk and Horstmann, 2004; Knott et al., 2008; Hayworth et al., 2014), and segmentation of 3D datasets (Saalfeld et al., 2010; Cardona et al., 2012; Meirovitch et al., 2016; Lee et al., 2017; Parag et al., 2018; Pavarino, 2020), resulting in terabytes or even petabytes of data (Lichtman et al., 2014). As sample volume size increases, new protocols must be developed and optimized including sample preparation, fixation, sectioning, and imaging. To meet all these needs, we have developed a pipeline for processing brain tissue from gastropod mollusc *Berghia stephanieae* (Valdés, 2005), which, for several reasons outlined below, has begun to attract considerable interest in neuroscience.

*Berghia* has features that make it amenable to a connectomic analysis of the neurons in its brain. Like other nudibranchs, the central ganglia are condensed in the head. The head ganglia have approximately 10,000 neurons in the adult. *Berghia* is smaller than other commonly used gastropods, being approximately 2 cm in body length. It has a short lifecycle with distinct developmental stages and can be easily cultured in the lab as is its food source, the anemone *Exaiptasia diaphana* Rapp, 1829.

Here, we outline several protocols developed for imaging *Berghia’s* brain using volume electron microscopy. These include high-pressure freezing and freeze substitution, a BROPA-based method (Mikula and Denk, 2015; Genoud et al., 2018) referred to here as ROPO (reduced-osmium-pyrogallol-osmium), and a rapid processing protocol. This pipeline could serve as a starting point for standardizing sample preparation protocol in other species.

## Materials and Methods

### Animal husbandry

Both *E. diaphana* and *B. stephanieae* were obtained from either ReefTown or Salty Underground and reared separately in 10-gallon tanks with an air pump. Abiotic factors were measured weekly. Salinity was measured with an Aichose Brix refractometer (Shenzhen Xindacheng Commercial and Trading Co., Ltd.) and kept to a density of 1.021. pH was measured using standard pH test strips and maintained at 8.0. At least once a week, *B. stephanieae* were fed pieces or whole *E. diaphana*. Water changes were performed once a week.

### Berghia stephanieae neuroanatomy

Gross morphology of *Berghia’s* brain can be examined under DIC and stained with DAPI (Figure 1A, B). This image shows 2 pairs of ganglia - cerebropleural (CP) and pedal (P) (Figure 1A). Each rhinophore has a single rhinophore ganglion (Rh) (Figure 1A). The eyes and statocysts are embedded in the ganglion sheath between the cerebropleural and pedal ganglia (Figure 1A).

**Figure 1.**
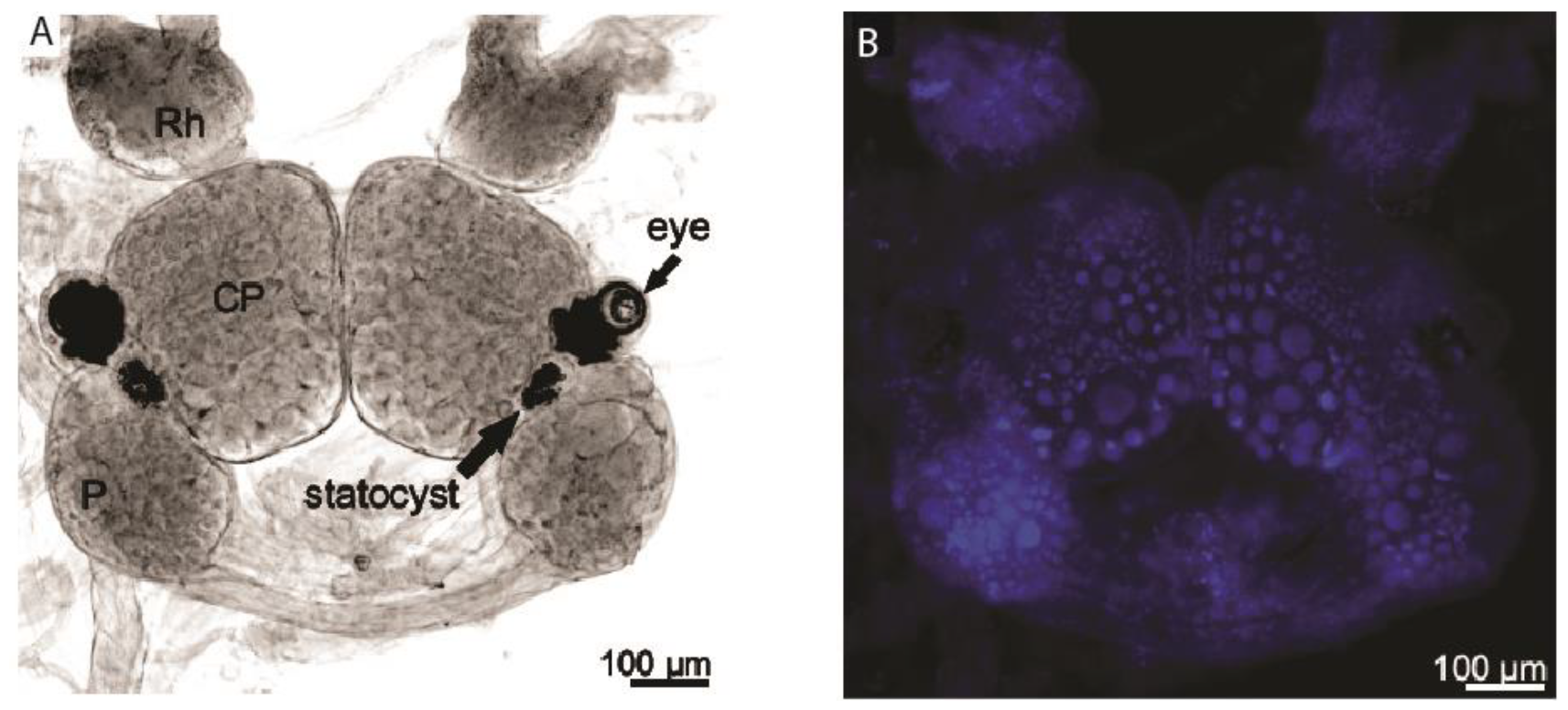
Adult brain of *Berghia stephanieae*. (A) Adult brains dissected and imaged under light microscopy reveal paired ganglia: rhinophore (Rh), cerebropleural (CP) and pedal (P). A pair of statocysts and eyes are connected to the brain. (B) Staining with DAPI shows how variable the cells and their nuclei are in size.

### Sample Processing Methods: HPF-FS, ROPO, and rapid processing protocol

We developed a pipeline for processing tissue, sectioning, and imaging *Berghia’s* brain (Figure 2). Whole brains were processed from adults with a body length of 0.8 – 1.0 cm. The brain, consisting of the cerebropleural, pedal, buccal, and rhinophore ganglia was quickly removed from the body using two forceps, one to hold the oral region near the buccal mass, and one to clamp behind the rhinophores and pulling the forceps apart.

**Figure 2.**
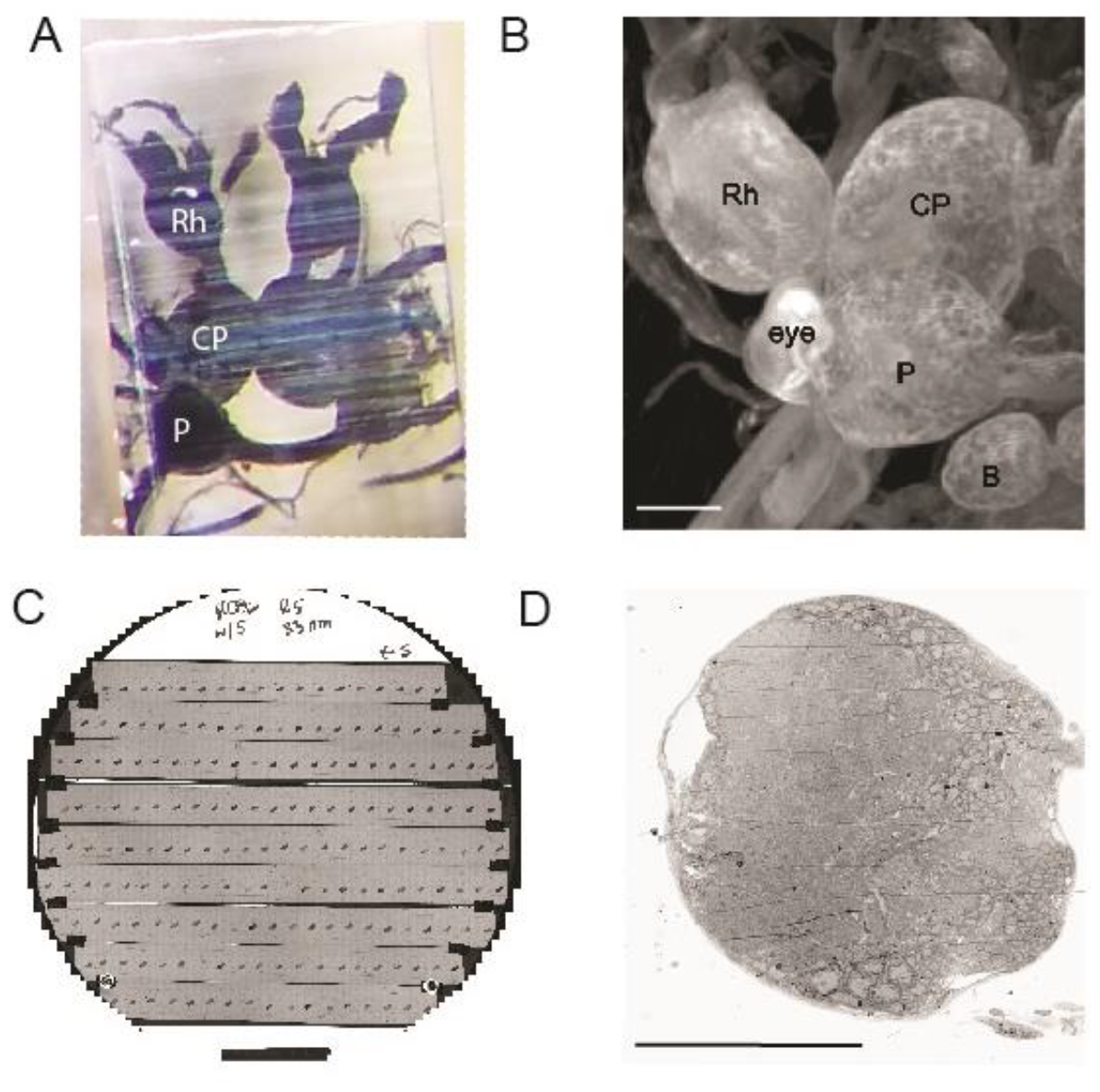
Pipeline for processing samples. (A) Adult brains are dissected and processed before being embedded and cured in resin. (B) µCT imaging provides higher resolution projections post-EM processing. All ganglia and eyes are present. It also determines orientation of the sample and distance from the cutting surface. (C) Image of a wafer with sections placed after serially sectioning the tissue in ATUM microtome. (D) A single slice of brain tissue after imaged sections are stitched together in the XY plane. All scale bars - 100µm. Rh - rhinophore ganglion, CP - Cerebropleural ganglion, P - Pedal ganglion, B - Buccal ganglion.

High-pressure freezing and freeze-substitution (HPF-FS) were performed at Harvard Medical School using the Leica EM ICE and EM AFS-2 (Leica Microsystems Inc, Buffalo Grove, IL). Whole brains were immediately frozen upon dissection. Excess water was removed from the holder with a Kimwipe or by loading the holder while on top of a piece of filter paper and sliding the holder into place for freezing. HPF was followed by 72-hour incubation in a solution of 1% glutaraldehyde, 1% OsO_4_, 0.1% tannic acid, 0.1% uranyl acetate (UA), and 1% dH_2_O in acetone. Samples were left in the solution for 48 hours at -90°C and gradually warmed to room temperature over 24 hours. After briefly rinsing samples in acetone, they were infiltrated in 1:1 mixture of acetone and LX112 resin (15.6 g LX112, 4.7 g NSA, 9.7 g NMA, 0.6 g BDMA; LADD research Industries, Williston, VT) for at least three hours, further infiltrated in full resin overnight, and cured in fresh resin in flat bottom capsules (Ted Pella, Inc., Redding, CA) at 60°C for 48 hours.

Tissue processed with the ROPO protocol was performed at Harvard with all steps performed at room temperature and on a rocker. Brains were dissected from adults (1 cm BL) in primary fixative consisting of 1% glutaraldehyde, 3% mannitol, 9% sucrose, and 0.4 M sodium cacodylate buffer (pH 7.8) and left in fixative for one hour in 20 mL scintillation vials. Samples were washed twice in buffer for 10 minutes followed by a one-hour incubation in 1.5% K_4_FeCN_6_/1% OsO_4_. After washing in dH_2_O twice, samples were incubated in 0.5 M pyrogallol for one hour and washed twice again in dH_2_O. At this point, samples were often transferred to a new holder due to precipitate formed between pyrogallol and osmium. After an hour incubation in 2% OsO_4_, samples were dehydrated in an ascending series of ethanol (50%, 70%, 95%, 2 × 100%, 10 minutes each) followed by acetone for 10 minutes. Samples were infiltrated and cured as above.

A rapid processing protocol was developed by taking the ROPO protocol and manually pipetting each solution, including the washes, over a sample at least ten times using fresh borosilicate glass pipets for each step. Infiltration in 1:1 acetone: resin was also performed manually by pipetting several times. Samples were allowed to sit in full resin on a rocker overnight before being cured as above.

### Sectioning and Imaging

Blocks were trimmed to a hexagonal block face approximately 1 × 2 mm with a 45° diamond trimming knife (DiATOME, Hatfield, PA) followed by imaging in a Zeiss Xradia 520 Versa 3D X-ray microscope (Carl Zeiss Microscopy LLC) at 40 kV and 3W for up to 30 hours. This was done to assess tissue quality, relative size of individual ganglia, and orientation with respect to the sectioning plane. Sections for screening tissue and serial sections were obtained on an automated tape ultramicrotome (ATUM; Hayworth et al., 2006; Schalek et al., 2012) and Leica EM UC6 ultramicrotome (Leica Microsystems, Buffalo Grove, IL) at 30-35 nm with 4 mm 35° diamond knives (DiATOME, Hatfield, PA). Reels of collected tape were cut into strips and attached to 100 mm diameter silicon wafers via double-sided conductive tape. If tissue was not *en bloc* stained, then after plasma cleaning wafers were stained with ∼10 mL of 4% UA for 4 minutes, rinsed twice in dH_2_O, and dried with pressurized air before repeating with ∼10 mL of 2% lead citrate for 4 minutes, rinsed twice, and dried. Only post-staining with lead citrate was performed for samples processed with HPF-FS. Fiducial markers were affixed to the double-sided tape at the corners of the wafer. Wafers were placed in a vacuumed chamber for at least 24 hours before imaging. Optical images for use in WaferMapper (Hayworth et al., 2014) were acquired on a Zeiss Axio with Axiocam 503 mono camera and Zen software (Carl Zeiss Microscopy LLC).

## Results

### High-pressure freezing and freeze-substitution

Samples are dissected and cryofixed in artificial seawater. The speed of brain dissection is crucial when processing with HPF-FS since one has only a few moments to transfer a dissected brain to the cryoholders and freeze. Once done, care must be taken to keep the holder and now cryopreserved sample under liquid nitrogen before freeze-substituting the nitrogen out for fixatives. Several freeze-substitution solutions of different concentrations and comprised of different chemicals (e.g., absence of UA) were tested. A major pitfall with FS is that samples show high electron density so discerning synapses versus membranes is difficult when processed in higher concentration than those reported here or were poorly preserved especially if water content was above 1%. The addition of tannic acid provided necessary contrast, but mitochondria and Golgi were nearly or in some cases completely electron opaque. Thus, while we emphasize the use of tannic acid, its concentration should be carefully considered. Subsequently, uranyl acetate was essential for membrane preservation and high contrast. Sections through cerebropleural ganglion (Figure 3A, B) showed well-preserved cells noted by the darkly stained membranes and nuclei. Similarly, sections through the neuropil of the rhinophore ganglion showed excellent preservation of processes and extracellular space (Figure 3C, D).

**Figure 3.**
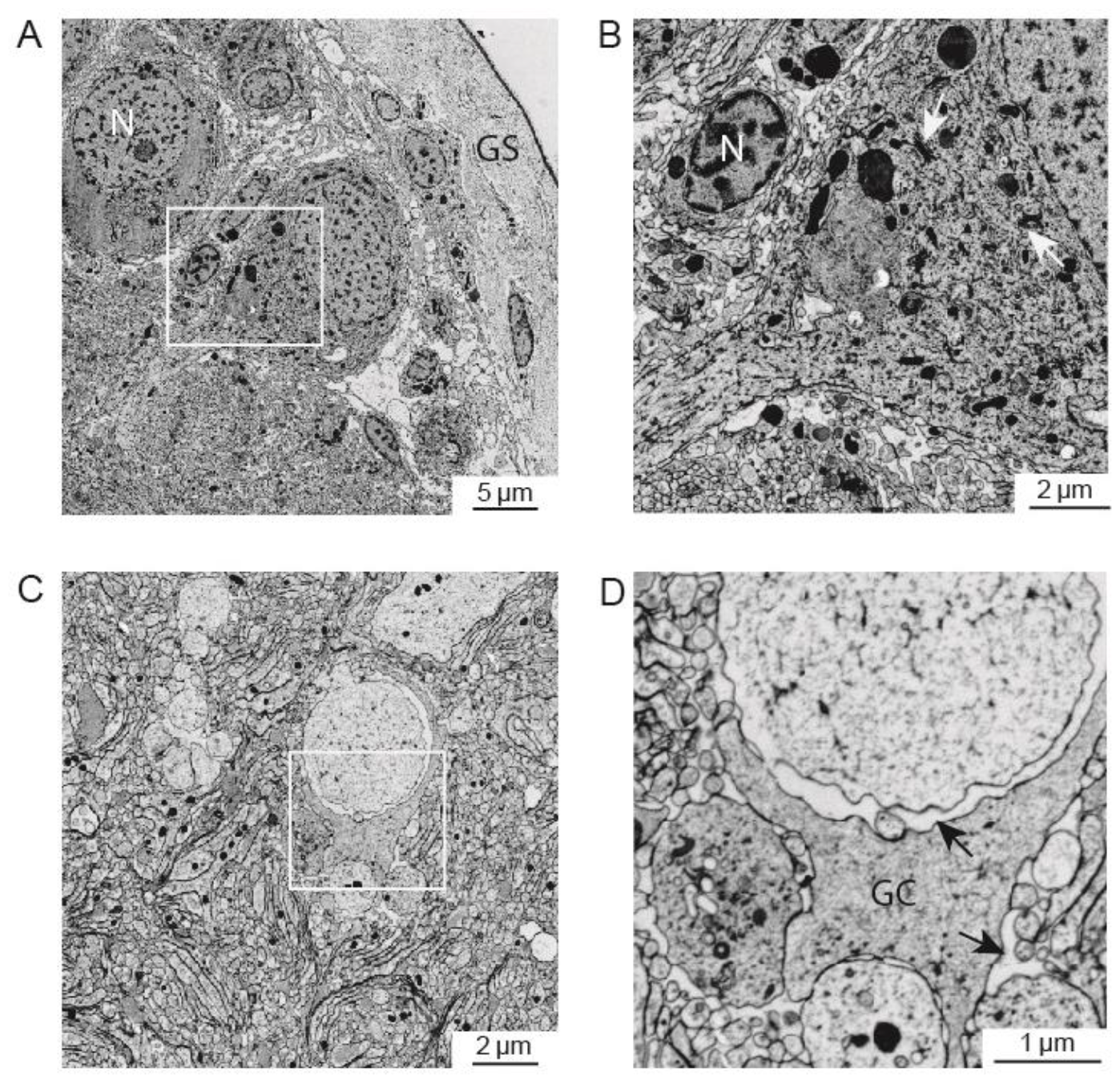
HPF-FS results of adult brain from *Berghia stephanieae*. (A) HPF-FS provided high contrast and preservation of the outermost ∼200 µm tissue in the cerebropleural ganglion including (B, box in A) the ganglion sheath (GS), nuclei (N) and golgi apparatus (arrows in B). (C,D) Section through the rhinophore ganglion shows excellent preservation of extracellular space (arrows) particularly around a glial cell (GC) clearly seen at higher magnification (D).

### Aldehyde-based tissue processing

One of the first issues to tackle with aldehyde-based methods was preservation of extracellular space that would allow traceability. The addition of mannitol into the primary fixative provided the closest likeness to samples processed via HPF-FS with enough extracellular space for tracing and segmentation. No other substitute tested (e.g., sucrose, cysteine) came close to mannitol in providing adequate fixation. Additionally, the level of contrast resulting from HPF-FS needed to be similar when using aldehyde-based methods especially since the former is limited to depth of cryopreservation. In addition, finding a repeatable and reproducible method of fixing and staining tissues was a major hurdle.

The result was the replacement of TCH in the common ROTO (reduced-osmium-thiocarbohydrazide-osmium; Hua et al., 2015) with pyrogallol (i.e., ROPO). Notable differences between ROTO and ROPO included fewer cracks or loss of tissue in the latter method (Figure 4A), and a more evenly distributed staining throughout the tissue (Figure 4B). When processed with TCH, ganglion sheaths were disrupted resulting in loss of peripheral cell bodies, torn connectives, and large holes in the tissue blocks, presumably from nitrogen production (Mikula and Denk, 2015), incomplete staining, or regions of high osmium deposition not suitable for segmentation (data not shown). Although using pyrogallol in place of TCH resulted in some areas of tissue with what appeared to be heavy osmium deposition in the extracellular space (Figure 4B), tracing was still possible. These regions made identifying cell body membranes easier, but not the information present outside the cells. In other regions of the same tissue, the contrast was not as heavy, and one could discern extracellular material between the cell bodies (Figure 4C). Additional or longer washes did not seem to mitigate this result.

**Figure 4.**
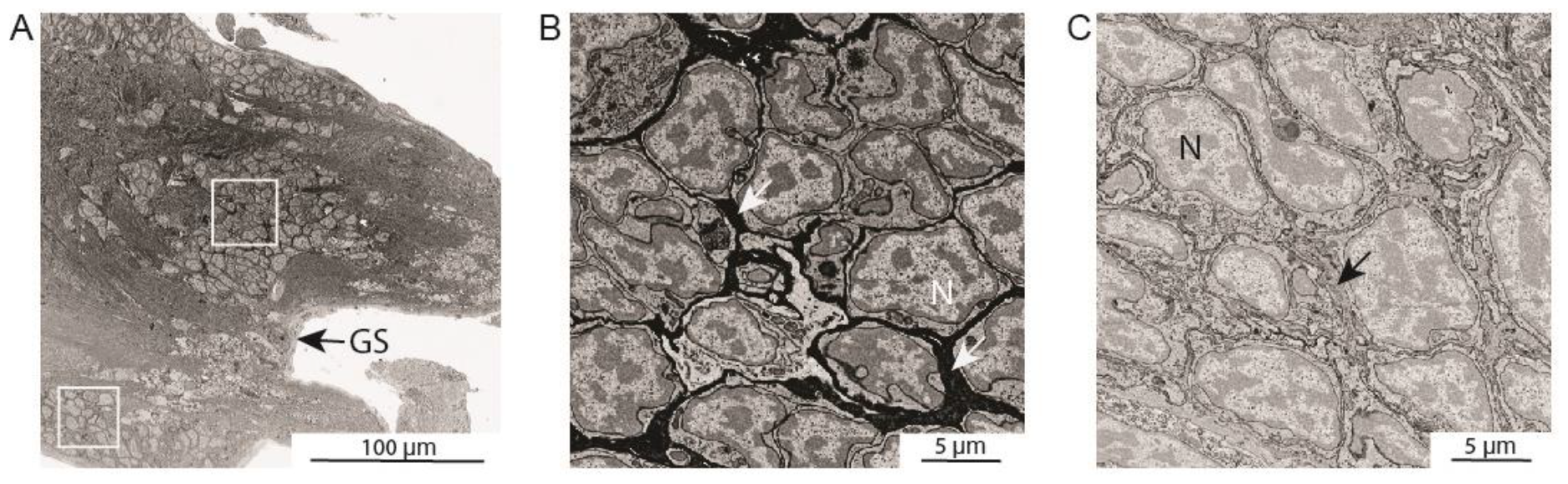
ROPO staining results in well preserved but variable electron density across tissue. (A) A section through the cerebropleural ganglion with well preserved ganglion sheath (GS) and cell bodies. (B) At higher resolution (from the top box in A) extracellular space is electron dense and surrounds cell bodies indicated by nuclei (N). (C) From the bottom box in A, extracellular space is not electron dense and material can be observed between cells.

Pyrogallol requires no heat or increased agitation to dissolve in water but using fresh pyrogallol reduced variability. The solution turns yellow with increasing intensity within 24 hours and presumably loses activity after several days. *Berghia’s* eye tissue seemed much better after the replacement of TCH (data not shown).

### Staining and Embedding

Staining wafers with UA and LC provide enough post-sectioning contrast, but it is time consuming and uses up a large volume of each solution (∼10 mL per solution per wafer) especially when one has thousands of sections to stain. As such, *en bloc* staining methods were first tested with UA since it was part of the FS solution following HPF. Interestingly, when dissolved in water UA formed a dark precipitate just beneath the ganglion sheath while nuclei remained unstained (Figure 5A). When UA was dissolved in acetone as it is in FS, no precipitate formed, and nuclei were adequately stained. However, due to the hazards and increasing regulation on acquiring UA depending on the country in which one conducts research (e.g., Japan and Canada), we found that gadolinium acetate and samarium acetate (Odriozola et al., 2017) were suitable alternatives to UA. Both gadolinium and samarium are non-radioactive lanthanides and while neither separately or together provided the high contrast typical of UA, they sufficiently stained nucleic acids (Figure 5C, D) and thus negated the need for UA. These alternatives were not tested with HPF-FS, but the results reported here from use in the ROPO method should be considered if higher contrast is needed and UA is not easily available. For enhancing protein and membrane contrast, copper sulfate lead citrate *en bloc* stain developed by Tapia et al. (2012) (Figure 5B) proved more reliable and reproducible than lead aspartate (not shown). In particular, the membranes of nuclei, mitochondria, and rough ER were well-preserved and opaque (Figure 5B) compared to unstained samples. It is important to remember that while one may have optimized staining, serial sectioning suffers from wrinkles and folds, and reducing these artifacts is heavily influenced by the choice of resin.

**Figure 5.**
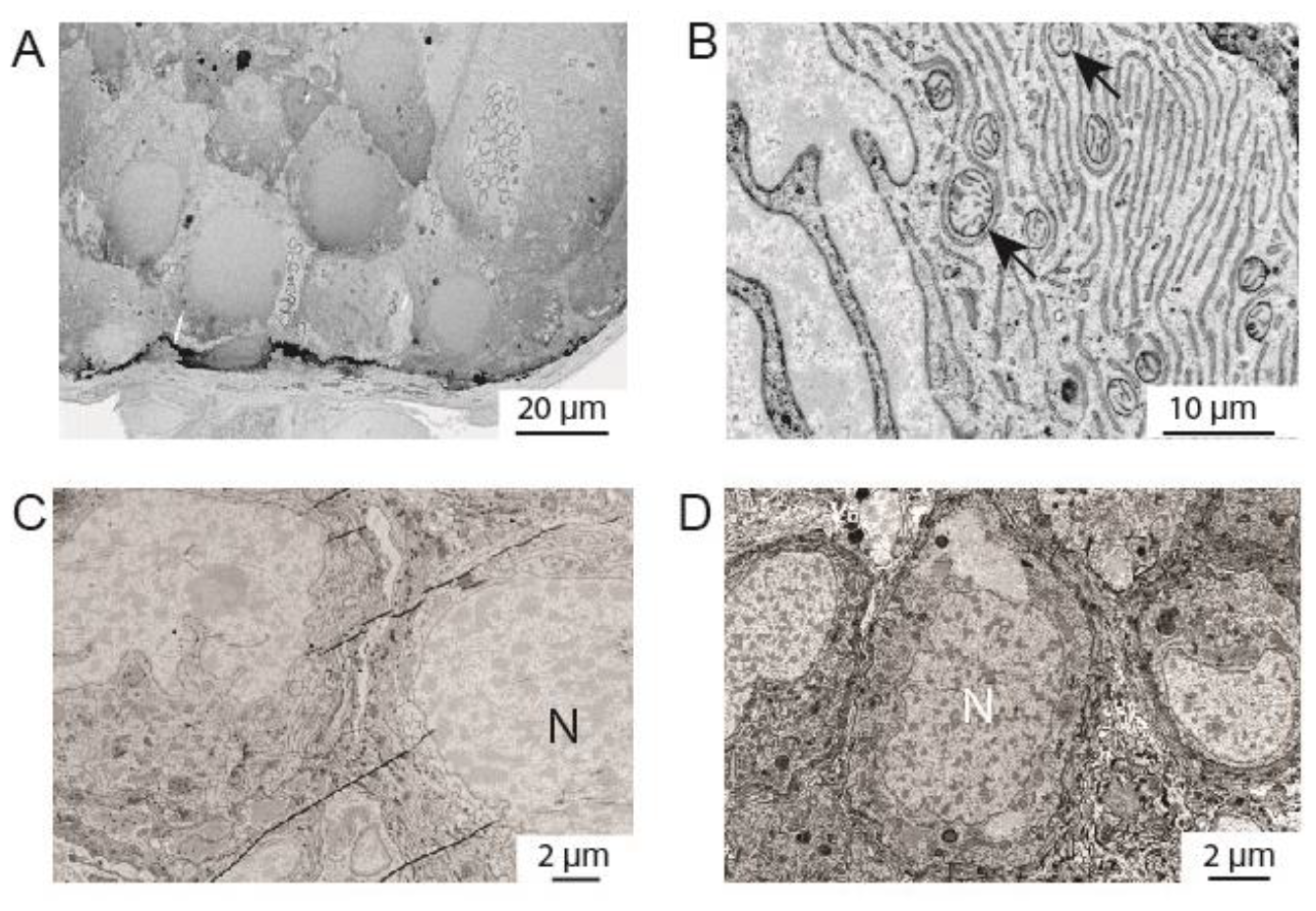
En bloc staining of brain tissue. (A) Section through cerebropleural ganglion with UA dissolved in water resulted in unstained nuclei (N) and a dark precipitate just beneath the ganglion sheath (arrows). (B) En bloc staining with copper sulfate lead citrate (CSLC) reliably stained cell membranes. Organelles like mitochondria (arrow) can also been seen. (C) Gadolinium acetate and (D) samarium acetate resulted in a well stained sample with distinct nuclei (N).

It was difficult to find a suitable resin due to the shape of the brain. The ganglia are fairly round in cross-section and the amount of space between the ganglia and their connectives resulted in high number of wrinkles. After many tests comparing resins such as EMBed812 (Epon), Durcupan, and Araldite, we chose the less viscous and harder LX112. Infiltration was superior and fewer wrinkles were observed compared to samples cured in the commonly used Epon. With the brain, eye, and juvenile successfully processed with new protocols, we went further to ascertain what other techniques might be applicable for preserving neural tissue, more specifically with respect to processing time.

### Rapid Processing

The ROPO method along with *en bloc* staining takes approximately two days usually with an overnight infiltration step. We developed a rapid processing protocol for adult brain tissue after learning about the mPrep system developed by Microscopy Innovations, LLC. The system is an automated processor that involves a constant flow of chemicals through the sample chamber, and by doing so processes entire samples in a fraction of the time conventional protocols require. Since there was little need for a machine dedicated to processing hundreds of samples and could not justify the cost, a more hands-on protocol was adapted. The results were promising for contrast and overall preservation, and there was no observable chemical or mechanical damage to the tissue (Figure 6A, B). The lack of extracellular space compared to that in non-rapid processing (Figure 6C) was an important issue for dissected brain samples. However, this is one avenue of potential interest for those looking to reduce hands-on time without the expense of automated machinery and brings one closer to generating a high-resolution brain atlas for connectomic analysis.

**Figure 6.**
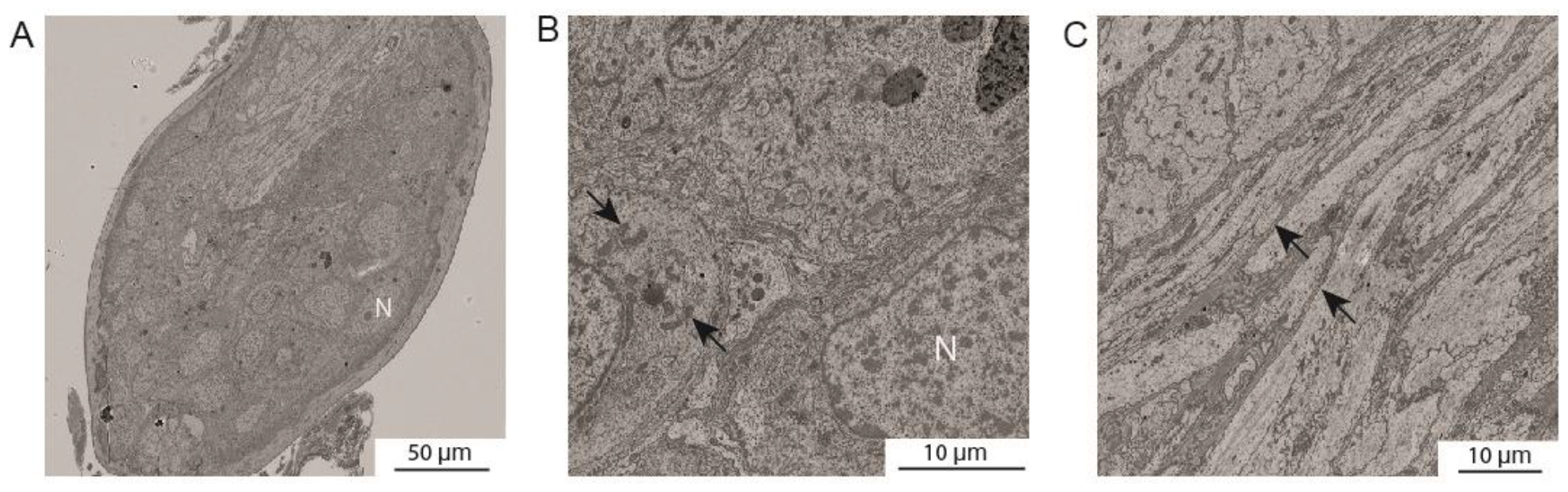
Rapid processing of brain tissue. (A) An oblique section through cerebropleural ganglion shows good contrast with no tissue damage from processing or sectioning. Higher-resolution images from dashed boxes in (A) showed good ultrastructural preservation of organelles including nuclei (N) and mitochondria (arrows in B), but lack of extracellular space (arrows in (C)).

## Discussion

We describe several methods used in a general EM processing pipeline for fixing, staining, and imaging brain tissue of gastropod mollusc *Berghia stephanieae*. The most repeatable method was HPF-FS. However, HPF reliably preserves tissue only up to 200 µm. Nonetheless, HPF-FS was instrumental in our development of the ROPO protocol. ROPO is repeatable and superior to ROTO protocols for our samples. However, pyrogallol results in an unusual staining pattern. The differences in contrast in the extracellular space within the same ganglia point to a possible difference in the biochemistry of these regions that attract a greater amount of osmium. We do not know enough about the biochemistry within the brain to understand why pyrogallol and osmium might be found in greater or lesser amounts across the tissue. The tissue also had some wrinkles and folds in different regions across sections. Machine and deep learning techniques are being developed on our results to correct for these artifacts.

Rapid processing protocol is promising and has a quick turnover in fixing and staining samples. A protocol that involves rapid processing and preserves extracellular space would be a highly advantageous step for other animals too.

Research for understanding neural circuitry in gastropod molluscs is being undertaken at an ever-growing pace now that the techniques for imaging and segmenting at synaptic resolution for this organism have been developed and deployed. This project also paves the way for a new era in comparative neuroanatomy, by providing the necessary methodology to facilitate large-volume connectomics and neural reconstructions in diverse organisms.

## Acknowledgements

We thank Bhoomi Patel and Nika Enringros for establishing developmental timelines of *Berghia*. Kelly Fischer and Daniela Molina-Palacios for their help in standardizing sample preparation protocols. Shuohong Wang and Yuelong Wu for their help in stitching sections.

## References

Berger, D. R., Seung, H. S., Lichtman, J. W. (2018). VAST (Volume Annotation and Segmentation Tool): efficient manual and semiautomatic labeling of large 3D image stacks. Front. Neural Circuit. 12, 88. 10.3389/fncir.2018.00088

Genoud, C., Titze, B., Graff-Meyer, A., Friedrich, R. W. (2018). Fast homogeneous en bloc staining of large tissue samples for volume electron microscopy. Front. Neuroanat. 12, 1–8.

Hayworth, K. J., Morgan, J. L., Schalek, R., Berger, D. R., Hildebrand, D. G. C., Lichtman, J. W. (2014). Imaging ATUM ultrathin section libraries with WaferMapper: a multi-scale approach to EM reconstruction of neural circuits. Front. Neural Circuit. 8, 1–18.

Hayworth, K. J., Kasthuri, N., Schalek, R., Lichtman, J. W. (2006). Automating the collection of ultrathin serial sections for large volume TEM reconstructions. Microsc. Microanal. 12(Supp 2), 86–87.

Hua, Y., Laserstein, P., Helmstaedter, M. (2015). Large-volume en bloc staining for electron microscopy-based connectomics. Nat. Commun. 6, 7923. 10.1038/ncomms8923

Kristof, A., Klussman-Kolb, A. (2010). Neuromuscular development of Aeolidiella stephanieae Valdéz, 2005 (Mollusca, Gastropoda, Nudibranchia). Front. Zool. 7, 5. 10.1186/1742-9994-7-5

Januszewski, M., Kornfeld, J., Li, P. H., Pope, A., Blakely, T., Lindsey, L., et al. (2018). High-precision automated reconstruction of neurons with flood-filling networks. Nat. Methods. 15, 605–610.

Meirovitch, Y., Matveev, A., Saribekyan, H., Budden, D., Rolnick, D., Odor, G., Knowles-Barley, S., Jones, T. R., Pfister, H., et al. (2016). A multi-pass approach to large-scale connectomics. https://arxiv.org/abs/1612.02120

Mikula, S., Denk, W. (2015). High-resolution whole-brain staining for electron microscopic circuit reconstruction. Nat. Methods. 12, 541–546.

Montanaro, J., Gruber, D., Leisch, N. (2016). Improved ultrastructure of marine invertebrates using non-toxic buffers. PeerJ. 4, e1860. 10.7717/peerj.1860

Odriozola, A., Llodrá, J., Radecke, J., Ruegsegger, C., Tschanz, S., Saxena, S., et al. (2017). High contrast staining for serial block face scanning electron microscopy without uranyl acetate. bioRxic [Preprint]. Available at: https://www.biorxiv.org/content/10.1101/207472v1 (Accessed July 23, 2019).

Parag, T., Tschopp, F., Grisaitis, W., Turaga, S. C., Zhang, X., Matejek, B., Kamentsky, L., Lichtman, J. W., Pfister, H. (2018). Anisotropic EM segmentation by 3D affinity learning and agglomeration. https://arxiv.org/abs/1707.08935

Pavarino, E., Berger, D. R., Morozova, O., Lichtman, J. W., Meirovitch, Y. (2019). mEMbrain: an interactive deep learning tool for labeling and instance segmentation of EM datasets. Berstein Conference 2019 Abstract. 10.12751/nncn.ba2019.0083.

Quinlan, P. D., Fischer, K. E., Drescher, B., Lichtman, J. W., Katz, P. S. (2019). Spatial vision from a low-resolution eye. Neuroscience Meeting Planner Program No. 067.08.

Society for Neuroscience, Chicago, IL. Ronneberger, O., Fischer, P., Brox, T. (2015). U-net: Convolutional networks for biomedical image segmentation. 1505.04597.

Saalfeld, S., Cardona, A., Hartenstein, V., Tomančák, P. (2010). As-rigid-as-possible mosaicking and serial section registration of large ssTEM datasets. Bioinformatics. 26, i57–i63.

Schalek, R., Kasthuri, N., Hayworth, K., Berger, D., Tapia, J., Morgan, J., Turaga, S., Fagerholm, E., Seung, H., Lichtman, J. (2011). Development of high-throughput, high-resolution 3D reconstruction of large-volume biological tissue using automated tape collection ultramicrotomy and scanning electron microscopy. Microsc. Microanal. 17(Supp 2), 966–967.

Lee, K., Zung, J., Li, P., Jain, V., Seung, H. S. (2017). Superhuman accuracy on the SNEMI3D connectomics challenge. 31^st^ Conference on Neural Information Processing Systems, Long Beach, CA, USA. https://arxiv.org/abs/1706.00120

